# Prevalence and Risk Factors Associated to Non-Communicable Diseases in Khartoum State

**DOI:** 10.1101/711028

**Authors:** Samah Mohamed Aabdin Sayed, Ayman Mohamed Aabdin, Mohammed Altyb Alshykh Abo-Shanab, Mounkaila Noma

## Abstract

**Introduction:** Non-communicable diseases are multifactorial including genetic, physiological, environmental behavioral factors. Our research aimed to determine the prevalence and risk factors associated with Non-communicable diseases in two administrative units of Khartoum State.

**Methods:** A community-based cross-sectional study was conducted in two administrative units of Khartoum State on a sample of 132 participants selected through multi-stage sampling technique. Firstly, a stratified random sampling technique was used to select Alshohada/Soba out of the six administrative units of Khartoum locality (mode of living was urban). In Jabaal Awliya locality of four administrative units, Al Jabal (with urban and rural mode of living) was selected. At second level, 50 households were selected in each of the two administrative units through the geographical information system to obtain a representative spatial distribution of households in each of the administrative areas. At third level, in each of the household selected participants experiencing at least one NCD were included in the study after obtaining his/her verbal well informed consent. The data collected were computerized through Epi Info 7 and analyzed through SPSS 23. The data were firstly summarized numerically and graphically. Association among variables were determined through chi-square tests and ANOVA. A multi-logistic regression was conducted to estimate the risk factors associated to NCDs. All statistical tests were considered significant when *p* < 0.05.

**Results:** Our findings revealed that NCDs prevailed with an overall prevalence of 24/100,000 population. Of the fifteen risk factors associated to NCDs in the two administrative units, seven were statistically associated (*p* < 0.05) to NCDs.

**Discussion:** In our research the risk factors statistically associated with NCDs were age and gender of the participants, their profession, educational level, physical activities, follow-up visits and having meals outside home. In conclusion NCDs were public health priorities with particular attention to diabetes and hypertension to avoid premature deaths.

## INTRODUCTION

Non-communicable diseases (NCDs), also named chronic diseases, are multifactorial including genetic, physiological, environmental and behavioral factors as reported by the World Health Organization [1]. WHO reported that globally 41 million deaths occurred yearly due to NCDs with 15 million of those losing life were aged 30 to 69 years; over 85% of these premature deaths occurred in low- and middle-income countries. The four leading killers accounting for 80% of all premature NCD deaths were cardiovascular diseases (17.9 million death/year), cancers (9.0 million), respiratory diseases (3.9 million), and diabetes (1.6 million). The risk of dying from NCDs increases with tobacco use, physical inactivity, the harmful use of alcohol and unhealthy diets. By reducing these risk factors, WHO challenged the participant countries to meet the sustainable development goal (SDG) related to non-communicable diseases and mental health by reducing by one third the premature NCD mortality [1].

Regarding the socioeconomic impact of NCDs, there is a close relation between poverty and NCDs. People at lower socioeconomic position get sicker and die earlier than people at higher social class. This is due to many reasons such as improper diet and poor access to health services [2]. A systematic analysis carried on age-sex specific mortality for 282 causes of death in 195 countries and territories revealed that non-communicable diseases, as a leading cause of deaths, contributed by 73.4% to the total deaths in 2017 [3].

In Sudan, the proportional mortality rate ranked cardiovascular disease as first accounting for 28% of the total deaths, follow up by cancers (6%), chronic respiratory diseases (3%) and diabetes (2%). In the overall, NCDs in Sudan accounted for 52% of all deaths. The risk of premature death between 30-70 years in Sudan was reported to 26% with a gender variation between males (28%) and females 24% [4].

A multi-country longitudinal study on aging and health [5] revealed a prevalence of hypertension was high in all six countries covered (China, Ghana, India, Mexico, Russia and South Africa). The self-reported prevalence of hypertension ranged from 14.1% in Ghana to 40.8% in Russia. Followed by self-reported arthritis with a prevalence ranging from 7% in Mexico to 27.2% in Russia. The prevalence of self-reported asthma was 7.7% in India, whereas in the remaining countries the prevalence was < 5%. The prevalence of both self-reported and algorithm based chronic lung diseases were lowest in Ghana 0.5% and 3% respectively, and highest in Russia 13.7% and 21.4% respectively. The adjusted prevalence of self-reported depression varied from 0.3% in China to 13.9% in Mexico. Whereas, the algorithm based indicated a low prevalence of depression in China (2.1%) and a high prevalence in India (18.5%).

Population-based cross sectional studies revealed that a prevalence of having at least one chronic disease of 45.1% in the age group of ≥ 18 years [6], while in adults aged ≥ 35 years the reported prevalence of NCDs of 22.8% varied from 22.0% in urban areas to 24.0% in rural setting [7]. In those aged ≥ 60 years, an overall prevalence of NCDs of 31.7% was published [8]. Of the four NCDs of concerned hypertension ranked first (26.0%), followed by diabetes mellitus (8.0%), stroke (1.9%) and chronic obstructive pulmonary disease (COPD, 1.0%).In India, NCDs accounted for 60% of all deaths which represented one third of the total deaths due to NCDs in South-East Asia Region (SEAR) of WHO with four NCDs taking the lead cardiovascular diseases (45%), chronic respiratory diseases (22%), cancer (12%) and diabetes (3%) as confirmed by an Indian author who reported a prevalence of NCDs of 29.0% with 31% suffering from chronic obstructive pulmonary disease, 29% from hypertension, 24% from chronic heart disease [9,10].

NCDs remained a challenge even in displaced populations. a 2015 study [11] on thirteen displaced camps in Kurdistan Region of Iraq involved 8360 participants, the results revealed one-third of households interviewed had at least one member who had one or more of the four common NCDs which are: hypertension (19.4%), musculoskeletal conditions (13.5%), DM (9.7%) and cardiovascular disease (CVD, 6.3%). In another cross-sectional household survey among 8041 non-camp Syrian refugees aged ≥ 18 years, Rehl et al. found that 21.8% of study population suffered from at least one NCD and 44.7% reported NCD multi-morbidities. Hypertension and diabetes prevalence were respectively 14.0% and 9.2% [12]. The prevalence of NCDs and use of health services was implemented in Gaza Strip on a sample of 760 households totalizing 5192 individuals; the findings revealed that 12.7% of the participants harbored at least one NCD [13].

The African continent also paid tribute to NCDs. WHO-Non communicable Diseases (NCDs) Country profiles of 2018 reported that for all deaths in Nigeria and Uganda 29 and 33 people out of 100 lost their lives because of NCDs [4]. A population-based cross-sectional survey using WHO STEPS survey instrument indicated that in Tanzania the prevalence of hypertension ranged from 16.0% to 17.0%, whereas in Uganda it ranged between 19.0% and 26.0%. In both countries hypertension was more prevalent in urban than in rural population. The prevalence of diabetes mellitus (DM) and other NCDs was low with a prevalence of 1 to 4% for DM [14, 15]. The Malawi National NCDI Poverty Commission reported that in 2017, diabetes, cardiovascular diseases, cancer, and chronic lung diseases were account for 38% of the burden of diseases from NCDs. These findings were consolidated in 2018 review which revealed that the prevalence of cardiovascular diseases was 8.9%, asthma prevalence was 5.0% and the prevalence of diabetes ranged between 2.3% and 5.7% and all types of cancers incidence rate was 156 cases/100,000 population [16,17]. In the Democratic Republic of Congo, Mawaw P M et al [18] reported through a cross-sectional study conducted among 2,749 employees the prevalent NCDs were hypertension (18.2%), diabetes (11.7%), obesity (4.5%) and CVD (3.2%). The facility-based retrospective study conducted in Ethiopia in 2015 on 22,320 medical records revealed that the reason for the visits due to NCDs represented 29.7% of the records. The prevalence of cardiovascular diseases was 18.8% and of diabetes mellitus 13.1% [19]. A systematic review [20] of five hospital-based studies from eastern Ethiopia revealed a prevalence of CVD was 7.2% and 2.4% for hypertensive heart diseases among all age groups. The only available study among adult outpatients of the capital city indicated that prevalence of CVD was 24%. Two studies investigated the prevalence of hypertensive heart diseases in the capital city, the findings revealed a prevalence of 12% among adults aged ≥11 years. The prevalence of diabetes was assessed through two studies one in the Southern region among adults aged ≥ 18 years and the second in urban region. Two additional hospital based studies, one in all age group and the second in patients aged ≥ 20 years revealed a prevalence of diabetes of respectively 0.5% and 1.2%. These two studies reported a prevalence of diabetes of respectively 4.9% and 5.3%. Regarding cancer, one hospital-based study on outpatient adults aged ≥ 20 years reported a prevalence of 0.3%. The prevalence of asthma was estimated through one community based study and two hospital-based studies patients aged ≥ 20 years. The three studies revealed a prevalence of asthma of respectively 0.6%, 1% and 3.5%.

In Sudan, published articles [21,22] revealed a prevalence of hypertension ranging from 27.6% to 35.7%. Abdalla EAM et al. reported that diabetes mellitus among 236 adults was 18.6% with no significant gender difference in the prevalence rate [23].

The risk factors associated to non-communicable disease are multifactorial. Ahmed Reza Hosseinpoor et al. reviewed 2002-04 WHS Data from the 41 countries that had available data about NCD risk factors, and relevant socioeconomic and demographic variables. Their findings revealed that angina, asthma, arthritis and depression prevalence were inversely associated to wealth and education, on the contrary the prevalence of diabetes showed a strong association with wealth and education [24].

A meta-analysis in Japan estimated the excess risks on deaths and life expectancy based on data from the National Health and Nutrition Survey and epidemiological studies. The findings revealed that tobacco smoking (129,000 deaths [95% CI: 115,000-154,000]), high blood pressure 104,000 deaths (95% CI: 86,000-119,000) and physical inactivity (52,000 deaths, [95% CI: 47,000-58,000]) ranked first [25].

A cross sectional study, using the WHO Stepwise approach for surveillance of NCDs, was implemented on sample of 3,489 participants (51.7% urban and 48.3) aged 15-64 years. Their findings revealed that the prevalence of daily smoking was 6.6% in urban and 12.29% in rural participants. The prevalence of low physical activity of 38.9% in urban participants was lower (14.2%) in rural; the prevalence of low fruit and vegetables consumption of 92.7% was higher (96.4%) in rural participants. Raised blood pressure of 29.1% in urbans was found to be 15.4% in rural participants [26]. Another cross-sectional study in Ibadan (Nigeria) using the WHO Stepwise approach [27] to assess the risk factors of NCDs among 606 civil servants indicated that the prevalence of smoking, harmful use of alcohol, low physical activity, insufficient fruit and vegetables intake, obesity were respectively 6.5%, 7.8%, 62.2%, 69.7%, 57.3%. In the same country, Agaba EL et al. implemented a survey of non-communicable diseases and their risk factors among university employees by using WHO Stepwise approach [28]. Of 883 participants of the University of Jos (Nigeria), the most common NCDs were hypertension (48.5%), chronic kidney disease (13.6%), and diabetes mellitus (8.0%). The most common NCD risk factors reported were inadequate intake of fruit and vegetables (94.6%), physical inactivity (77.8%), obesity (26.7%), alcohol use (24.0%) and cigarette smoking (2.9%). In Uganda, a community-based cross-sectional study using WHO stepwise approach to chronic disease risk factors surveillance revealed on a sample of 518 participants, that 20.5% of females and 22.1% of males had hypertension, 9.0% of participants were diabetics, 4.9% of men and 9.0% of women were obese, and 51% of participants were physically inactive [29].

In Sudan, Ghebreselasie D.T. [30] reported during the 2nd annual congress of medicare expo on primary care and general pediatrics, that NCDs accounted for 44.0% of the overall deaths in Sudan. Based on a sample 380 participants aged ≥ 30 years selected through a stratified random sampling technique he pointed out that a prevalence of smoking in males of 18.4% and 0.3% in females; alcohol consumption in males was 3.9%, physical inactivity in both gender was 75.0% and adequate consumption of fruits and vegetables was prevalent in 72.9% of the study population. History of NCDs was statistically associated (*p* < 0.05) to cigarette/tobacco use, as well as physical activity and educational level with a *p-value* of respectively 0.003 and 0.011.

Our study attempted determine the prevalence and risk factors associated with non-communicable diseases in two administrative units (Alshohada/Soba and Al Jabal) of Khartoum State.

### Methodology

A community-based cross-sectional study was conducted. The research was implemented in two administrative units of Khartoum State, namely Alshohada & Soba and Al Jabal with a total population of 552,955 people, 51.1% living in Alshohada & Soba and 48.9% in Al Jabal as per Sudan Census Bureau of Statistics (http://cbs.gov.sd/). The population of Alshohada/ Soba is distributed in 33,608 households and 270,662 inhabitants of Al Jabal live in a total of 32,537 households. A multistage sampling technique was used to select the study participants. At first level the two localities (administrative level 2) which administratively constitute Khartoum were all included in the study. Al Khartoum locality comprise six administrative units, one administrative unit (Alshohada & Soba) was randomly selected. In the locality of Jabal Awliya, subdivided in four administrative units, the only administrative unit where the population mode of living is rural and urban, Al Jabal was purposely included in the study to enable to have in the total sample both urban and rural setting. At second level, in each of the two select administrative units, 50 households spatially distributed were included in the study under the assumption that in every one of the households selected one case of NCD will be available. When no case of NCD was found in a selected household, it was systematically replaced by the nearest household. This led to an estimated sample size of at least 100 participants (50 households x 2 administrative units).

Inclusion and exclusion criteria. Were included in the study, the residents of Alshohada/Soba and Al Jabal harboring a non-communicable disease, regardless their gender and age; for the participants below < 15 years the questionnaire was addressed by the care giver present at the time of the data collection. Were excluded from the all residents from other administrative units of Khartoum State and the residents of the two selected administrative units who refused to participate in the study.

A standardized interviewer-administrated questionnaire, pre-tested, primarily developed in English, was administrated in Arabic by the researcher. The data collected included the socio-demographic characteristics of the study participants (age, gender, residence, occupation, household size), the type of NCD, its duration, comorbidity and factors associated to the NCD including the lifestyle of the participants and the management of the NCD.

The data collected were computerized through a template developed in Epi-Info7 and analyzed through the statistical package for social sciences (SPSS 23). The data were summarized numerically (mean, standard deviation, median) and graphically (frequency tables for estimating prevalence and graphics). Association between categorical variables were through chi-square tests (Pearson Chi-square, Fischer Exact Test and Likelihood Ratio). A multi-logistic regression analysis was performed to assess the relationship between NCDs and its related risk factors. All statistical tests were considered as statistically significant when *p* < 0.05.

## RESULTS

### Characteristics of the study participants

Our research was conducted in two administrative units of Khartoum State. 54.5% (72/132) of the participants lived in Alshohada & Soba and the remaining 45.5% (60/132) where from Al Jabal. Females were predominant (55.3%, 73/132) than males (44.7%, 59/132). The age of the participants ranged from 4 to 90 years with a median age of 56.5 years. 70.8% (92/132) of the participants were married; 44.7% (59/132) were housewives, 37.9% (50/132) were working and 17.4% (32/132) were not. 86.2% (112/130) were educated and the remaining 13.8% (18/130) did not attend a school. Their respective household size varied from 2 to 14 members with a median of 6 members.

### Distribution of non-communicable diseases (NCDs) in two localities of Khartoum State Chronic Diseases reported by the study population

The two most frequent NCDs reported were Diabetes mellitus and Hypertension affecting respectively 33.0% (44/132) and 23.5% (31/132) of the participants. 21.1% (28/132) of the participants harbored the two NCDs (diabetes mellitus associated to hypertension). 22.1% (29/132) of the participants suffered from a single NCD or combined two to three NCDs.

### Prevalence of non-communicable diseases Alshohada-Soba and Al Jabal

The overall prevalence of the NCDs in the two administrative units surveyed was 24/100,000 population. It ranged from 22/100,000 population in Al Jabal to 26/100,000 population in Al shohada-Soba.

Diabetes Mellitus was the most prevalent (8/100,000) NCD in the two administrative units. This prevalence was the same in Al shohada-Soba and Al Jabal with a prevalence of respectively 7.79/100,000 and 8.13/100,000 (table 1).

**Table 1:** Prevalence (case/100000 population) of NCDs in Alshohada & Soba and Al Jabal (n=132)

| Non-communicable disease | Number and prevalence of NCDs per administrative unit |  |  |  |  |  |
| --- | --- | --- | --- | --- | --- | --- |
|  | Alshohada-Soba | Prev* | Al Jabal | Prev | Total | Prev |
| Diabetes mellitus | 22 | 7.79 | 22 | 8.13 | 44 | 7.96 |
| Hypertension | 16 | 5.67 | 15 | 5.54 | 31 | 5.61 |
| Diabetes & Hypertension | 19 | 6.73 | 9 | 3.33 | 28 | 5.06 |
| Asthma | 2 | 0.71 | 6 | 2.22 | 8 | 1.45 |
| Diabetes mellitus &<br>Cardiovascular disease | 4 | 1.42 | 0 | 0.00 | 4 | 0.72 |
| Diabetes mellitus &<br>Cardiovascular disease &<br>Hypertension | 2 | 0.71 | 2 | 0.74 | 4 | 0.72 |
| Cardiovascular disease | 2 | 0.71 | 1 | 0.37 | 3 | 0.54 |
| Osteoarthritis | 0 | 0.00 | 3 | 1.11 | 3 | 0.54 |
| Cancer | 1 | 0.35 | 1 | 0.37 | 2 | 0.36 |
| Rheumatoid Arthritis | 1 | 0.35 | 1 | 0.37 | 2 | 0.36 |
| Hyperthyroidism | 1 | 0.35 | 0 | 0.00 | 1 | 0.18 |
| Hypothyroidism | 1 | 0.35 | 0 | 0.00 | 1 | 0.18 |
| Cardiovascular disease &<br>Hypertension | 1 | 0.35 | 0 | 0.00 | 1 | 0.18 |
| **Total** | **72** | **26** | **60** | **22** | **132** | **24** |

\*Prevalence calculated as number of cases of NCDs divided by 100,000 total populations.

Hypertension ranked second in term of prevalence, it affected 6 persons per 100,000 populations. This burden was equal in the two administrative units.

Twenty-eight participants reported harboring diabetes mellitus and hypertension, this represented an average of 5 per 100,000 total populations. This average varied across the two administrative units with 7 persons affected out of 100,000 populations in Alshohada-Soba and 3 persons/100,000 in Al Jabal.

### Risk factors associated with Non-Communicable Diseases

#### Practices of the study participants towards treatment and follow-up for NCDs

##### Adherence to treatment by the study participants

Of 132 participants, the majority (93.9%, 124/132) replied that they were under treatment, the proportion of participants under treatment was higher (97.2%, 70/124) in Alshohada & Soba than in Al Jabal (90.0%, 54/124). This difference between the two administrative units was not statistical significant *(p*=0.14). Most (72.6%, 90/124) of the participants were under treatment for ≥ 5 years and only 8.1% (10/124) were under treatment for < 1 year. 19.4% (24/124) of those were under treatment between 1-4 years. The participants were asked to provide the type of treatment they were using. The question offered three options which were namely medical, traditional or combined treatment. 91.9% (114/124) of the participants were under medical treatment and the remaining 8.1% (10/124) were under combined treatment (medical and traditional). In Alshohada & Soba, 54.4% and 80.0% of the participants were respectively under medical and combined treatment; whereas in Al Jabal those under medical treatment or combined were respectively 45.6% and 20.0%. The differences recorded between the two administrative units and the type of treatment were not statistically significant (*p*=0.107); this indicated that the type of treatment was equally used in both administrative units.

Anti-diabetic and hypertensive drugs were the most frequently used by the participants with respectively 63.7% (79/124) and 51.6% (64/124). 19.4% (24/124) of the participants were under other medications such as chemotherapy, neomercazol, statin, salbutamol, thyroxin, warfarin, and vitamins.

The participants were asked to address the question related to the adherence to treatment. Firstly 52.4% (65/124) have forgotten at least once to take his/her treatment and 47.6% (59/124) had never forgotten to take their treatment. Secondarily, 76.6% (95/124) of the participants were taking always their treatment.

##### Adherence to follow up as prescribed by the treating doctor

The participants were asked if “they have a regular follow up visit for their condition by their treating doctor”. More than half (52.3%, 69/132) always had regular visit, 9.8% (13/132) never visit their treating doctor for following their condition and 37.9% (50/132) visited their treating doctor only when necessary (Only when feel the need and only when suffered from the condition).

Sixty participants provided their reasons for not having regular follow up. The predominant (53.3%, 32/60) reason was the participant assumed that the follow-up visit was not a necessity, followed up by “being too busy” with 16.7% (10/60). Financial constraints and others (including long waiting time to see the doctor, home follow-up) ranked equally third with 15.0% (9/60).

##### Physical activities of the study participants

Of the 132 participants, 37.1% (49/132) reported to practice physical exercises with walking as the predominate exercise. 62.9% (83/132) did not practice any physical exercise. In working place, 60.8% (31/51) of the participant sat for > 2 hours at work and 55.3% (73/132) sat at home for > 2 hours. 64.4% (85/132) of the participants had an unhealthy sitting time either at work or at home.

In each of the two administrative units, the proportion of participants with an unhealthy sitting time at work/home was higher (70.8% in Alshohada-Soba vs 56.7% in Al Jabal) than those who had a healthy sitting time (29.2% vs 43.3%). However, there was no statistically significant association (*p*= 0.091) between administrative unit and time spent in sitting at workplace/home. A statistically significant association (*p*= 0.028) was found between the sitting time spent at workplace/home and gender.

### Dietary habits of the study participants

#### Food intakes

Fruit daily consumption was reported as to be “always” by 43.2% (57/132) of the participants, 54.5% (72/132) consumed fruits “occasionally” and 2.3% (3/132) “never” had fruits. In the overall fruits was consumed “always” by 43.2% of the participants and “occasionally/never” by 56.8%. The daily consumption of vegetables was reported as “always” by 65.9% (87/132) on the participants, 33.3% (44/132) had vegetables “occasionally” and one participant (0.8%) never ate vegetables. In conclusion, the daily consumption of vegetables was “always” for 65.9% of the participants and “occasionally/never” for the remaining 34.1% (45/132) participants.

The food intakes in a week were recorded through four variables which were namely having assida/kisra (meal made with sorghum/ millet/ maize), rice, red meat and salty foods (processed foods). Each of those variables was recorded as ‘always”, “Once a week/occasionally” and “never”. Regarding assida/kisra, most (70.5%, 93/132) of the participants had it “once a week/occasionally”, 18.9% (25/132) ate always assida/kisra, and 10.6% (14/132) never ate assida/kisra. Rice was consumed once a week by 83.3% (110/132) of the participants, 7.6% (10/132) daily and 9.1% (12/132) never had rice.

The frequency of regular (“always) consumption of salty foods in a week was higher (71.4%, 20/28) in Al Jabal than in Alshohada-Soba (28.6%, 8/28) where 60.2% (59/98) “never” consumed processed salty foods. The frequency of consumption of salty foods was statistically different (*p*=0.004) between the administrative units. The frequency of the foods was not statistically associated (*p* > 0.05) with the gender of the participants.

Having meals outside home in a week was recorded as “always”, “occasionally” for the participants who replied either once a week or occasionally and “never”. Having meals occasionally was reported by 69.7% (92/132) of the participants, 10.6% (14/132) had meals outside and 19.7% (26/132) had never ate outside home in a week. Of the 26 participants who reported never eat outside home, 50.0% lived in Alshohada-Soba and the remaining 50% resided in Al Jabal. The observed differences between frequency of having meals outside home and the administrative area of residence was not statistically significant (*p*= 0.158). The chi-square for trend of 2.638 indicated a not statistically difference (*p*=0.267) between gender and having meals outside.

Soft drink consumption collected as “always”, “once a week”, “occasionally” and “never” was recorded as healthy soft drink habit when participants answered “never” and those who replied either “once a week”, “occasionally” or “always” were recorded as having unhealthy habits towards soft drinks. 50.8% (67/132) reported having a healthy habit towards soft drinks.

Healthy juice drink habits were compared with unhealthy juice drink behavior. Participants who replied “drinking always juice without sugar” and those who answered that they “never drink juice with sugar” were coded as having a healthy juice drinking habits and those who answered by any other response to drinking juice without sugar or with sugar had their replies coded as an unhealthy juice drinking habits. In the overall, the majority (85.6%, 113/132) of the participants had an unhealthy habit towards drinking juice and only 14 participants out of hundred (19/132) had a healthy habit.

Regarding consumption of sweets, cakes and ice cream in a week, 78.8% (104/132) had an unhealthy habit and 21.2% (28/132) had healthy habits towards sweets. 86.4% reported to consume tea, 60.6% took coffee, smokers of cigarettes at the time of interview represented 5.3% and 1.5% reported that they used to have alcohol.

#### Relationship between non-communicable diseases and their associated factors

The NCDs reported by the study participants were regrouped in five categories namely diabetes mellitus (35.2%, 43/122), hypertension (23.0%, 28/122), diabetes mellitus associated to hypertension (23.0%, 28/122), cardiovascular disease associated to diabetes mellitus and hypertension (9.8%, 12/122) and other NCDs (9.0%, 11/122). The other NCDs included asthma, osteoarthritis, cancer, rheumatoid arthritis, hyperthyroidism and hypothyroidism. A logistic regression analysis was performed to estimate the risk factors associated to the NCDs above listed and the reference group in the model was the other NCDs.

Diabetes mellitus, table 2a revealed that none of the fifteen risk factors was statistically associated *(p* > 0.05) to the condition. However, physical activity highly contributed to the model by 8.5 times ([95 % CI: 1.002-71.903], *p*=0.050) as well as having meals outside home (OR=4.643, [95 % CI: 0.497-43.376], *p*=0.178), time of sitting at workplace/home (OR=3.691, [95 % CI: 0.544-25.038], *p*=0.181) and adherence to follow up of medical visits (OR=2.405, [95 % CI: 0.573-10.084], *p*=0.230). Consumption of fruits contributed to the model by 1.8 times ([95 % CI: 0.260-12.419], *p*= 0.552) and marital status by 1.6 times ([95 % CI: 0.411-6.538], *p*= 0.483). Other risk factors contributing to the model for > 1 time to the model were consumption of vegetables, administrative unit of residence and education as indicated by table 2a.

**Table 2a:** Logistic regression estimating the fifteen risk factors associated to four NCDs

| NCDs | Risk factor | B | Wald | df | p-value | OR | 95% CI for OR |  |
| --- | --- | --- | --- | --- | --- | --- | --- | --- |
|  |  |  |  |  |  |  | Lower | Upper |
| Diabetes Mellitus | Intercept | -2.372 | 0.209 | 1 | 0.648 |  |  |  |
|  | Administrative unit | 0.210 | 0.060 | 1 | 0.806 | 1.234 | 0.230 | 6.619 |
|  | Gender | -1.447 | 1.919 | 1 | 0.166 | 0.235 | 0.030 | 1.823 |
|  | Age in years | 0.023 | 0.567 | 1 | 0.451 | 1.023 | 0.964 | 1.086 |
|  | Education | 0.321 | 0.335 | 1 | 0.562 | 1.378 | 0.465 | 4.082 |
|  | Marital Status | 0.495 | 0.492 | 1 | 0.483 | 1.640 | 0.411 | 6.538 |
|  | Profession | -1.282 | 2.301 | 1 | 0.129 | 0.278 | 0.053 | 1.454 |
|  | Household size | -0.097 | 0.266 | 1 | 0.606 | 0.908 | 0.628 | 1.311 |
|  | Treatment duration (years) | -0.600 | 0.428 | 1 | 0.513 | 0.549 | 0.091 | 3.317 |
|  | Adherence to treatment | -0.956 | 0.856 | 1 | 0.355 | 0.384 | 0.051 | 2.911 |
|  | Follow up visits | 0.877 | 1.439 | 1 | 0.230 | 2.405 | 0.573 | 10.084 |
|  | Physical activities | 2.139 | 3.849 | 1 | 0.050 | 8.488 | 1.002 | 71.903 |
|  | Vegetables | 0.170 | 0.021 | 1 | 0.884 | 1.185 | 0.121 | 11.570 |
|  | Fruits | 0.587 | 0.355 | 1 | 0.552 | 1.799 | 0.260 | 12.419 |
|  | Meals outside home | 1.535 | 1.813 | 1 | 0.178 | 4.643 | 0.497 | 43.376 |
|  | Sitting (home/work place) | 1.306 | 1.788 | 1 | 0.181 | 3.691 | 0.544 | 25.038 |
| Hypertension | Intercept | -3.098 | 0.279 | 1 | 0.598 |  |  |  |
|  | Administrative unit | 0.092 | 0.009 | 1 | 0.923 | 1.096 | 0.172 | 6.982 |
|  | Gender | -2.489 | 3.493 | 1 | 0.062 | 0.083 | 0.006 | 1.129 |
|  | Age in years | 0.071 | 3.855 | 1 | 0.050 | 1.074 | 1.000 | 1.152 |
|  | Education | 0.023 | 0.001 | 1 | 0.971 | 1.023 | 0.297 | 3.524 |
|  | Marital Status | 0.648 | 0.712 | 1 | 0.399 | 1.911 | 0.424 | 8.608 |
|  | Profession | -2.306 | 5.135 | 1 | 0.023 | 0.100 | 0.014 | 0.732 |
|  | Household size | -0.141 | 0.478 | 1 | 0.489 | 0.868 | 0.582 | 1.296 |
|  | Treatment duration (years) | -0.700 | 0.478 | 1 | 0.489 | 0.497 | 0.068 | 3.607 |
|  | Adherence to treatment | -0.493 | 0.200 | 1 | 0.655 | 0.611 | 0.071 | 5.289 |
|  | Follow up visits | 1.583 | 4.076 | 1 | 0.044 | 4.871 | 1.047 | 22.658 |
|  | Physical activities | 2.757 | 5.365 | 1 | 0.021 | 15.760 | 1.528 | 162.522 |
|  | Vegetables | 0.977 | 0.648 | 1 | 0.421 | 2.657 | 0.246 | 28.668 |
|  | Fruits | 0.642 | 0.359 | 1 | 0.549 | 1.900 | 0.233 | 15.516 |
|  | Meals outside home | 1.603 | 1.788 | 1 | 0.181 | 4.969 | 0.474 | 52.105 |
|  | Sitting (home/work place) | 0.562 | 0.290 | 1 | 0.590 | 1.754 | 0.227 | 13.565 |

Hypertension, a statistically significant association was found between hypertension and physical activity which contributed by 16 times ([95 % CI: 1.528-162.522], *p=* 0.021); other statistically significant risk factors were profession (*p*=0.023) and adherence to follow up visit (*p*=0.044) contributing to explain the status of hypertension in the study population by respectively < 1 time [95% CI: 0.014-0.732] and 5 times [95% CI: 1.047-22.658]. Despite a not statistically significant association with hypertension, having meals outside home contributed to explain the hypertension by 5 times ([95% CI: 0.474-52.105], *p=*0.181) and consuming vegetables contribute by 3 times ([95% CI: 0.246-28.668], *p*=0.421). Education, age in years, administrative unit, time of sitting at home/work place, consuming fruits, marital status contributed to model by more than 1 time.

Diabetes associated to hypertension, the two conditions were statistically associated with profession and education with a *p-value* of respectively 0.001 and 0.045. Despite a not statistically significant association, with an increasing contribution of respectively 6.8 to 9.9 times, physical activity (OR=6.830, [95% CI: 0.622-74.932], *p*= 0.116) treatment duration, consumption of vegetables and having meals outside home (OR=9.873, [95% CI: 0.824-118.305], *p*= 0.071) explained the association of diabetes and hypertension. Whereas, the time of sitting at home/workplace contribute by 4 times (OR: 3.984, 95% CI: 0.488-32.509], *p*=0.197). Table 2b displayed the other risk factors contributing for < 2 times.

**Table 2b:**
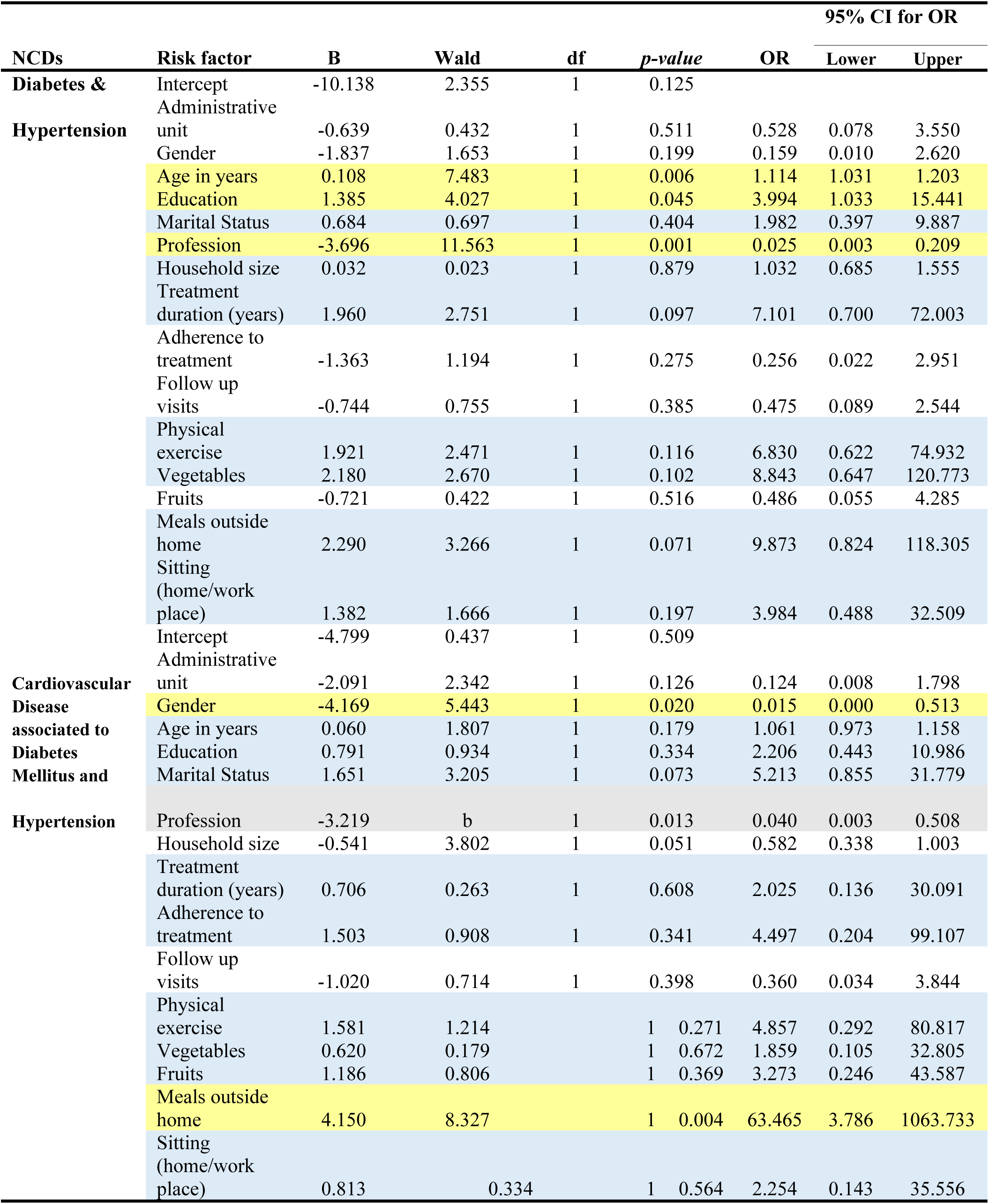
Logistic regression estimating the fifteen risk factors associated to four NCDs

Cardiovascular Disease associated to diabetes mellitus and Hypertension; the highly statistically significant contributor associated to this three conditions was having meals outside home contributing for 63 times ([95% CI: 3.786-1063.733], *p*=0.004). Another statistically significant risk factor was gender (*p*=0.020) and its coefficient of contribution of −4.169 indicated that males might be more likely to have the association of the three conditions. Table 2b revealed an increasing contribution ranging from 2 times to 5.2 times for treatment duration, education, time of sitting (home/work place), fruit consumption, adherence to treatment, physical activity and marital status.

In conclusion, in our research the risk factors statistically associated with NCDs were age, gender of the participants, their profession, education level, physical activities, follow-up visits and having meals outside home (Tables 2a and 2b).

## DISCUSSION

Our research sample of 132 participants was distributed in two administrative units (Alshohada-Soba and Al Jabal) of Khartoum Locality, they were aged 56.5 years (range: 4-90); females were predominant (55.3%) than males (44.7%).

The most frequent NCDs reported were diabetes mellitus (33.0%), hypertension (23.5%) and the comorbidity diabetes mellitus associated to hypertension (21.1%, 28/132). 22.1% of the participants suffered from a single NCD or combined two to three NCDs. In the overall, diabetes mellitus prevailed in 8 participants/100,000 total population, hypertension in 6 per 100,000 and the association diabetes mellitus affected 5 persons/100,000 populations. Abdalla EAM et al. [23] revealed in Jabra (Khartoum State, Sudan) a diabetes prevalence of 18.6% lower than ours, elsewhere in the literature in South Africa [31] a prevalence of diabetes of 5.0% was reported, in Kenya the prevalence of diabetes mellitus was 2.6% [32] and in Ethiopia, it ranged between 0.5% and 5.3% [20]. In South America, diabetes mellitus prevalence ranged from < 5.0% in Peru to 8.39% in Mexico [33,34]. Ma D. et al. revealed that diabetes prevalence was 6.9% in Japan, 9.2% in Korea and 13.1% in China [35].

Regarding hypertension, our prevalence of 23.5% was lower than those already reported by various authors who found a prevalence ranging from 27.6% to 35.7% [21,22,30]. On the African continent, a prevalence of 16.5% was reported in South Africa, and 24.8% in Kenya; whereas in South America, a hypertension prevalence of 11.14% was published in Mexico and 15.0%, in Peru. In Asia, the prevalence of hypertension was higher in China (24.5%) than in Korea (17.6%) and in Japan where a prevalence of 15.2% were published [31-35].

Ahmed M.H. et al. [36] reported a prevalence of 5.44% in 496 participants living with diabetes; in our research, cardiovascular disease was associated with diabetes mellitus in 3.0% of our study population. The combination of more than two NCDs was investigated by Alcalde-Rabanal JE et al. [34] in Mexico who reported the complex scenario of obesity, diabetes and hypertension on a sample of 10,326 participants and revealed that 73.0% suffered from diabetes, hypertension and/or overweight/obesity.

A statistically significant association between hours spent sitting at workplace/home and gender (*p*=0.028) and having salty food (processed food) and residence area (*p*=0.004).

A limitation of our research could be related to the use of a researcher designed questionnaire instead of using and adapting the WHO stepwise approach to chronic disease risk factor surveillance tool. Nonetheless, the research instrument developed provided data which through a multi logistic regression analysis enabled us to estimate the risk factors associated to one or more NCDs.

Our research indicated that the risk factors statistically associated with NCDs were age (OR= 1.114, 95% CI: 1.031-1.203], *p*= 0.006) in participants suffering from diabetes associated to hypertension, their gender (OR= 0.015, 95% CI: 0.000-0.513], *p*= 0.020) in participants suffering from the comorbidities cardiovascular disease, hypertension and diabetes, profession in patients living with hypertension (OR= 0.100, 95% CI: 0.014-0.732], *p*= 0.023) and those with diabetes associated to hypertension (OR= 0.025, 95% CI: 0.003-0.209], *p*=0.001). Education was statistically associated with the combined association of diabetes mellitus and hypertension (OR= 3.994, 95% CI: 1.033-15.441], *p*= 0.045). Having a regular follow up visit was statistically associated with the hypertension (OR= 4.871, 95% CI: 1.047-22.658], p= 0.044). Physical activities were statistically associated (OR= 15.76, 95% CI: 1.528-162.522], p= 0.021) in patients with hypertension. Having meals outside home (OR= 63.465, 95% CI: 3.786-1063.733], *p*= 0.004) was statistically associated and highly contributed with the combination of hypertension, diabetes and cardiovascular disease.

Non-communicable diseases are the results of complex interactions between the genetic make-up of an individual, lifestyle and environmental factors. In Sudan, age, gender, ethnic group, education level, family history of hypertension, family history of diabetes, residence, obesity, smoking, physical activity, salt and sugar intakes, renal problems, and pancreatic disease have been reported as risk factors of NCDs by various authors [21-23,30,36]. In the neighboring Ethiopia, low fruit consumption was reported [37] to account for 11.9% of NCD deaths which occurred in 2013, other risk factors published were alcohol consumption [31,33].

## Conclusions

Our research revealed that NCDs prevailed in Al shohada-Soba and Al Jabal with an overall prevalence of 24/100,000 populations. They were more frequent in Al shohada-Soba where 26 people out 100,000 were affected compared to Al Jabal where the prevalence was 22/100,000 populations. Of the fifteen risk factors associated to NCDs in the two administrative units, seven were statistically associated (*p* < 0.05) with NCDs.

NCDs should be a public health priority with particular attention to diabetes and hypertension which both well managed will prevent early deaths. Prevention and control of NCDs appeal collaborative efforts and sustained partnership between health services and the communities affected backed up by a strong political will and engagement to enable Sudan to implement WHO strategy towards reducing by one third the premature deaths related to NCDs by 2030.

## Acknowledgment

We would like to recognize and acknowledge our study participants whose contributions were crucial to bring our research to a successful end. Our sincere thanks to Mayada Abdarahman Abdin, Israa Giha, Sara Gabralla, Sara Kheiralla for their support and guidance throughout this research

## Supportive information

**S1:** Figure 1: Sampling frame of households in Soba-Alshohada delineated through Google Earth Pro 7.1.8.3036 (32-bit).

**S2:** Figure 2: geographical distributions of 50 households selected in Soba-Ashohada through Google Earth Pro 7.1.8.3036 (32-bit).

**S3:** Figure 3: Sampling frame of households in Aljabal delineated through Google Earth Pro 7.1.8.3036 (32-bit).

**S4:** Figure 4: geographical distributions of 50 households selected in Aljabal through Google Earth Pro 7.1.8.3036 (32-bit).

## Contribution of the Authors

**SAMS:** Designed, implemented the research, conducted the statistical analysis and drafted the initial manuscript.

**AMAS:** Actively in field implementation of the research and in computerizing the study data.

**MAAA:** Actively involved in field data collection and in data screening.

**MN:** Approved the research proposal, guided the statistical analysis and proofread the initial and the final manuscript.

**All the authors** read and approved the final version of the manuscript prior its submission.

## Data Availability

In accordance to data sharing and PLOS ONE policy on the matter, the authors declared that if the submitted manuscript is accepted for publication, the data will be deposit in the generalized repository of Dryad.

## Source of financing

The research was fully supported by Samah Mohamed Aabdin Sayed in the frame work of the fulfilment of her Master of Sciences in Public and Tropical Health in the University of Medical Sciences and Technology.

## Ethical clearance

The research proposal was approved by Sumasri Institutional Review Board in the framework of a community based survey. Authorization for the implementation of the research was obtained from each study participants after providing their well informed consent. The confidentiality of the participants was ensured through the use of anonymous research tool. The participants were reassured that the data collected from them will not be used for any other purpose other than the objectives assigned.

